# Light-induced condensates show accumulation-prone and less dynamic properties in the nucleus compared to the cytoplasm

**DOI:** 10.1101/2023.06.07.544154

**Authors:** Yuta Hamada, Akira Kitamura

## Abstract

Biomolecular condensates, including membraneless organelles, are ubiquitously observed in subcellular compartments. However, the accumulation and dynamic properties of arbitrarily induced condensates remain elusive. Here, we show the size, amount, and dynamic properties of subcellular condensates using various fluorescence spectroscopic imaging analyses. Spatial image correlation spectroscopy showed that the size of blue-light-induced condensates of cryptochrome 2-derived oligomerization tag (CRY2olig) tagged with a red fluorescent protein in the nucleus was not different from that in the cytoplasm. Fluorescence intensity measurements showed that the condensates in the nucleus were more prone to accumulation than those in the cytoplasm. Single-particle tracking analysis showed that the condensates in the nucleus are predisposed to be stationary dynamics compared to those in the cytoplasm. Therefore, the subcellular compartment may, in part, affect the characteristics of self-recruitment of biomolecules in the condensates and their movement property.

## 1. Introduction

Membraneless intracellular assemblies in subcellular compartments of cells are called biomolecular condensates [1]. Condensates are formed from spontaneous partitioning of biomolecules into discrete compartments of higher concentration compared to the surrounding solution through liquid-liquid and liquid-solid phase separation (LLPS and LSPS, respectively) [2]. In eukaryotic cells, various condensates containing protein and RNA, such as nucleolus, nuclear bodies, and nuclear speckles in the nucleus and processing bodies (PBs) and stress granules (SGs) in the cytoplasm, play diverse roles in cellular function and regulation of biochemical reactions [3]. For example, the nucleolus, a large multiphase liquid condensate and a famous membraneless organelle in the nucleus, is the basis for ribosome biogenesis and acts as a quality control compartment that contributes to protein homeostasis (proteostasis) [4]. PBs and SGs are condensates containing protein and RNA in the cytoplasm, but their functions are completely different: PBs are involved in mRNA turnover, and SGs are translational repressors during stress. [5] These differences in the function of condensates are currently known to be defined by the biomolecule composition in the condensates [5,6]. The interaction of the constituent biomolecules in the condensates is a controlling factor for the penetration of other molecules into the condensates [7]. As a simple and pure question, why do PBs and SGs not form in the nucleus? Alternatively, why do nuclear bodies and nuclear speckles not form in the cytoplasm? These questions prompted us to investigate whether the accumulation and dynamics of artificially formed intracellular condensates derived from the same protein differ between the cytoplasm and nucleus. To investigate this issue, we focused on the property of an *Arabidopsis thaliana* photoreceptor cryptochrome 2 (CRY2)-derived light-induced oligomerization tag, CRY2olig, that can form arbitrary clusters/condensates in cells [8,9]. Although wild-type CRY2 also forms intracellular assemblies by irradiation in plant and animal cells [10], CRY2olig carries mutations that are efficiently and reversibly oligomerized by blue light irradiation [9]. CRY2olig was tagged with a red fluorescent protein (mCherry) (hereafter called CRY2R); then, we compared the size, fluorescence intensity, and dynamics of the CRY2R condensates between the cytoplasm and nucleus using spatial image correlation spectroscopy (SICS) [11,12], fluorescence intensity measurement, and single-particle tracking analysis [13] based on confocal fluorescence microscopy.

## 2. Materials and Methods

### Cell culture and transfection

Neuro-2a murine neuroblastoma (N2a) cells were obtained from the American Type Culture Collection (ATCC; Manassas, VA, USA) and maintained in DMEM (08459-64, Nacalai Tesque, Kyoto, Japan) supplemented with 10% heat-inactivated fetal bovine serum (FBS) (12676029, Thermo Fisher Scientific, Waltham, MA, USA), 100 U/ml penicillin G (Sigma−Aldrich, St. Louis, MO, USA) and 0.1 mg/ml streptomycin (Sigma−Aldrich) as previously described [14]. One day before transfection, 2.0 × 10^5^ N2a cells were transferred to a glass bottom dish (3970-035, IWAKI-AGC Technoglass, Shizuoka, Japan). Plasmid DNA for expressing CRY2olig tagged with mCherry at its C-terminus was obtained from Addgene (#60032; Watertown, MA, USA). The plasmid (1.0 µg/dish) was transfected using Lipofectamine 2000 (Thermo Fisher Scientific) according to the manufacturer’s protocol. After incubation for 24 h, the medium was exchanged with normal growth medium before confocal observation.

### Light-induced oligomerization using a confocal microscope

Cell selection for observation, blue light irradiation, and image acquisition were performed using an inverted fluorescence microscope, Axioobserver Z1 (Carl Zeiss, Jena, Germany) combined with an LSM 510META (Carl Zeiss), a C-Apochromat 40x/1.2NA W Korr. UV−VIS-IR water immersion objective, and a heat incubator stage at 37°C and with 5% CO_2_. Cells expressing CRY2R were selected by ocular observation using fluorescence filter sets for mCherry (#49008, Chroma, Bellows Falls, VT, USA) with a mercury lamp. mCherry was excited at 594 nm (3.0 µW). Excitation and emission lights were split using a beam splitter (NFT488/594). Fluorescence was collected through a 615 nm long-pass filter (LP615) and then through a 545 nm dichroic mirror (NFT545). The diameter of the pinhole was 150 µm. After observation of 1 frame, the whole area of the cells was irradiated at 488 nm (0.66 µW); 1 iteration. After 488 nm irradiation, CRY2R-expressing cells were observed at 4.0 s intervals. The fluorescence intensity in the cells was measured 184–244 s after 488 nm irradiation when the condensates formed the most efficiently in the cells.

### Spatial image correlation spectroscopy

Spatial image correlation spectroscopy (SICS) analysis was performed as previously reported [12] from the confocal fluorescence images of CRY2R in N2a cells. Nonlinear curve fitting analysis for the distribution of the spatial autocorrelation function was performed on Octave software, including the calculation of the spatial autocorrelation function and its region cropping. The cytoplasmic and nuclear regions were distinguished by manually setting masks.

### Fluorescence intensity measurement

Fluorescence intensity in and out of the CRY2R condensate region was manually measured using Fiji-ImageJ (https://fiji.sc/). The mean intensities in CRY2R condensates in the cytoplasm and nucleus were normalized to those in the nucleoplasm or cytoplasmic region that did not include the condensates.

### Single-particle tracking

CRY2R condensates were induced in the same procedure as other measurements described above, and time series of confocal fluorescence images of the condensate-containing cells were acquired. The image series were analyzed using Fiji-ImageJ and the TrackMate plug-in [13]. CRY2R condensate particles were tracked in the cytoplasm and nucleus, and the total distance and mean speed of each particle were obtained.

### Statistics

The Student’s t tests were calculated using Microsoft-Excel.

## 3. Results

Accumulated clusters of CRY2R were induced by blue light irradiation in live N2a cells (Fig. 1a); however, it is still ambiguous whether the formed clusters are just oligomers or higher-order assemblies (i.e., condensates). To clarify this, we determined the mean size of the cluster using SICS, which can simply determine the average diameter of the cytoplasmic clusters [12]. The average diameter of the clusters both in the cytoplasm and in the nucleus (0.67–0.72 µm; Fig. 1b) was larger than the scale of the CRY2R oligomers. The diameter of the CRY2R clusters in the nucleus was not different from that in the cytoplasm (Fig. 1b). This suggests that the clusters may include many CRY2R molecules and can be defined as condensates in both the nucleus and cytoplasm.

**Figure 1.**
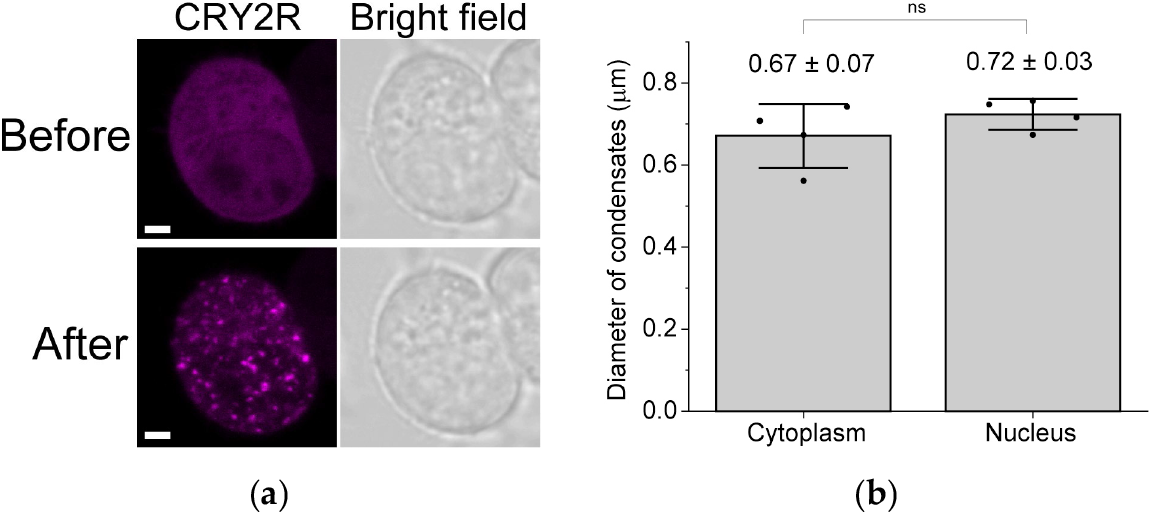
Size determination of CRY2R condensates using spatial image correlation spectroscopy (SICS). (**a**) Confocal fluorescence and bright field images of a typical N2a cell expressing CRY2R before and after blue light irradiation. Bar = 2 µm. (**b**) Box-and-whisker plot of the average diameter of CRY2R condensates in a single cell using SICS. The numerical values above the bars indicate the mean ± SD (n = 4). The *p* value was obtained from Student’s t test (ns: not significant; *p* <0.05).

Next, to investigate the extent to which molecules accumulate inside the condensates after blue light irradiation, the fluorescence intensity of CRY2R was compared in the condensates between the cytoplasm and the nucleus. The internal fluorescence intensity of the CRY2R condensates normalized using the intensity of the region that did not form the condensates was higher in the nucleus than in the cytoplasm (Fig. 2, a & b). The fluorescence intensity of the outside region of the condensates in the nucleoplasm was lower than that in the cytoplasm (Fig. 2c), likely because CRY2R rapidly accumulates in the condensates and the CRY2R concentration in the nucleoplasm decreases, but the CRY2R in the cytoplasm would not be immediately supplied to the nucleoplasm. These results suggest that CRY2R condensates in the nucleus may be more efficient in recruiting CRY2R outside the condensates than those in the cytoplasm.

**Figure 2.**
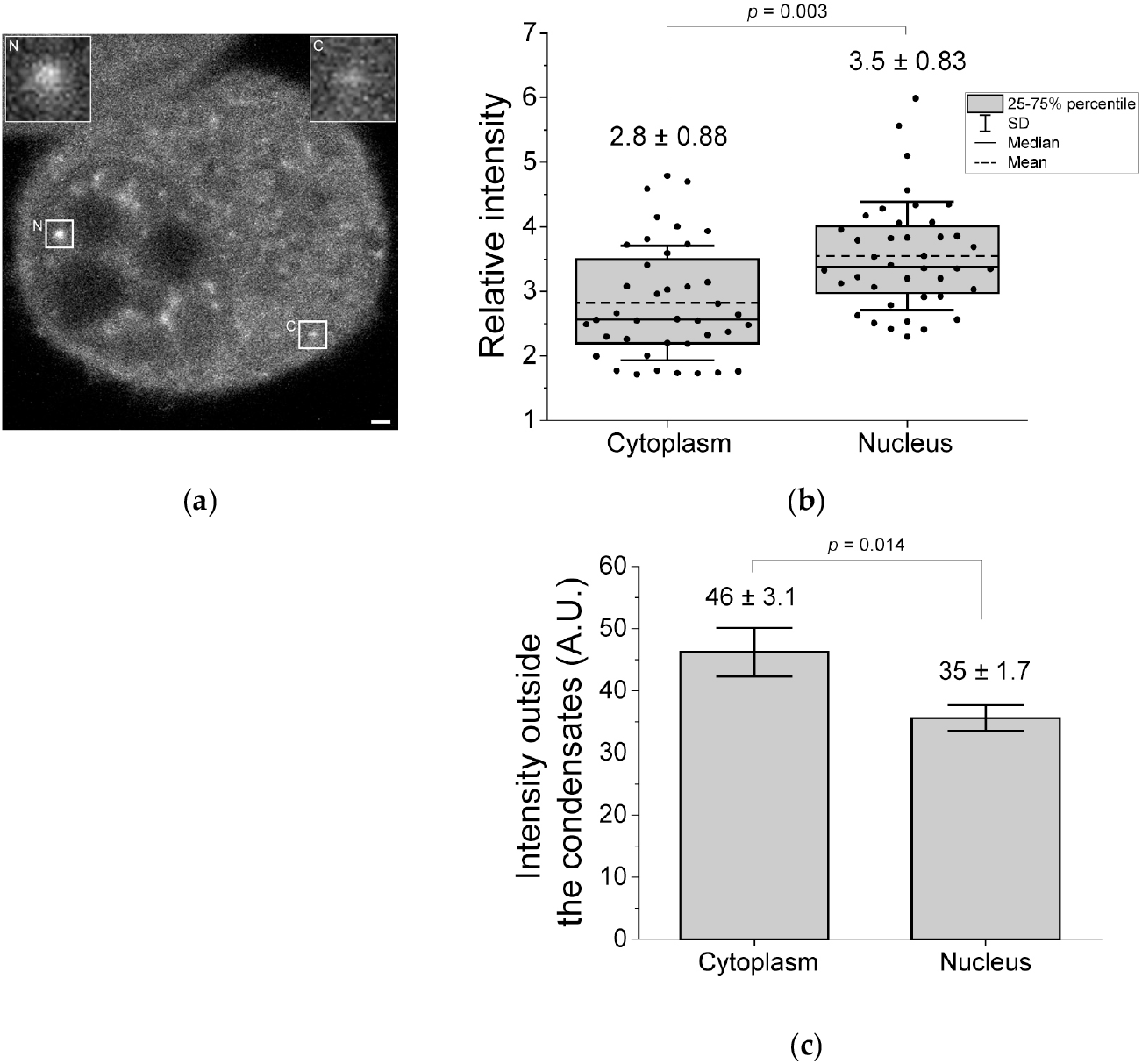
Comparison of the fluorescence intensity of CRY2R in condensates between the cytoplasm and the nucleus. (**a**) Confocal fluorescence image of CRY2R in live N2a cells. Typical condensates in the nucleus (N) and cytoplasm (C) are surrounded by white squares and enlarged (insets at the top). Bar = 1 µm. The images were interpolated using a bicubic interpolation algorithm. (**b**) The box-and-whisker plot of the relative intensity of the condensates to the outside region. The numerical values above the bars indicate the mean ± SD (40 condensates). (**c**) Bar plot of the fluorescence intensity of the outside region of the condensates used in (**b**). The numerical values above the bars indicate the mean ± SD (n = 3). The *p* value was obtained from Student’s t test.

Next, the dynamics (movement) of the CRY2R condensates in live N2a cells were compared between the cytoplasm and the nucleus (Videos S1 & S2). The mean speed of movement and the total distance traveled of the CRY2R condensates considering single particles during the observation period. The mean speed of movement and the total traveled distance of the CRY2R condensates were faster and longer in the cytoplasm than in the nucleus, respectively (Fig. 3, a & b). This indicates that CRY2R condensates in the cytoplasm tend to move around after formation, whereas those in the nucleus tend to stay where they are formed (i.e., stationary condensates). Furthermore, the mean speed of the cytoplasmic condensates varied, indicating that some of them were stationary, similar to those in the nucleus (Fig. 3b).

**Figure 3.**
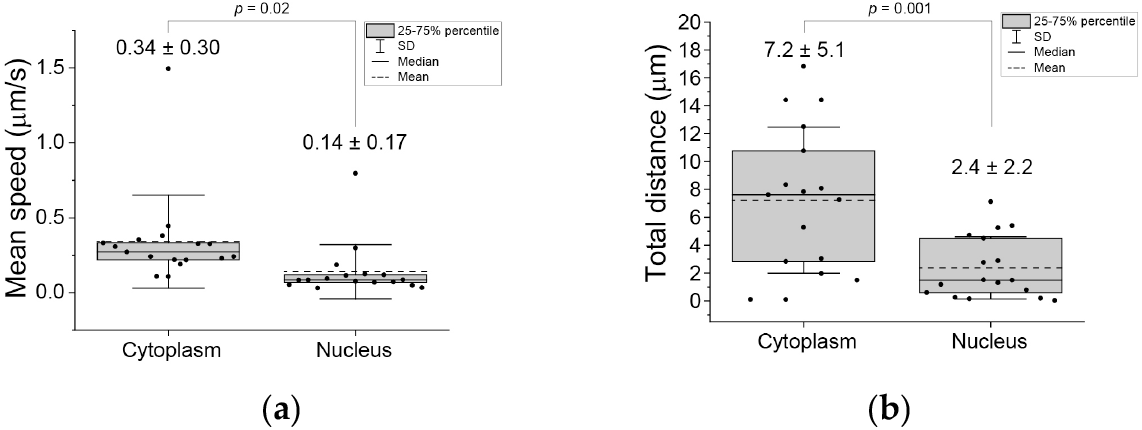
Single-particle tracking analysis of CRY2R condensates in live N2a cells. (**a**) Box-and-whisker plot of the mean speed (µm/s) of a single particle during image acquisition. (**b**) Box-and-whisker plot of the total traveled distance (µm) of single particles during image acquisition. The numerical values above the bars indicate the mean ± SD (17 particles). The *p* value was obtained from Student’s t tests.

## 4. Discussion

We compared the size, intensity, and dynamics of CRY2R condensates using confocal fluorescence microscopy and its application methods, such as SICS and single-particle tracking, in live cells. The size of CRY2R condensates determined using SICS (Fig. 1b) corresponds to the previously reported size [9]. This reaffirms the advantage of SICS, which is not good at precisely classifying objects/structures with multicomponent sizes but still provides an average value that is of equal significance. Endogenous *Arabidopsis* CRY2, the original full-length protein of CRY2olig, forms tetramers after blue light irradiation (∼12 nm in diameter) [15]. The size of the CRY2R condensates in the N2a cells was much larger than the determined oligomer structure; thus, they cannot be called ‘oligomers’, even considering the blur by optical diffraction. The fluorescence intensity ratio inside and outside of the condensates basically reflects the number of accumulated fluorescent molecules (Fig. 2b); however, since fluorescence lifetime of mCherry is shortened and quenched when it forms aggregates such as amyloids [16], this ratio may be less than the actual amounts of CRY2R in the condensates. Therefore, although oligomerization is probably an impetus for such condensates, these brightly accumulated structures in the cell could be called condensates, not just tetramers.

We showed that CRY2R formed stationary and highly accumulated condensates in the nucleus compared to those in the cytoplasm (Figs. 2 & 3). The lower concentration of CRY2R in the nucleoplasm than in the cytoplasm after the formation of the condensate suggests that the condensate in the nucleus likely appears to recruit CRY2R outside of the condensates (Fig. 2c), although a precise kinetic challenge remains. Nuclear localization of endogenous cryptochromes is conserved in animals and plants [17], suggesting the photoreceptor function of cryptochromes in the nucleus. An important interaction partner of cryptochrome is phytochrome, which is also localized in the nucleus [10]. However, since phytochrome is not expressed in N2a cells, what proteins contribute to the interaction with CRY2R? Since serine/arginine-rich splicing factor SC35 is colocalized with the blue light-induced condensates of CRY2olig in the nuclear speckles [9], SC35 is a candidate for the interaction that gives the stationary dynamics of CRY2R condensates. Viscosity and density in the subcellular compartment are assumed to be factors that make it prevention for the biomolecules to move. However, the microviscosity in the nucleoplasm does not change from that in the cytoplasm [18], and the density in the nucleoplasm is lower than that in the cytoplasm [19]; hence, neither the density nor viscosity in the subcellular compartment would be involved in the stationarity of the CRY2R condensates. Therefore, it is unquestionable that the interaction between CRY2R and endogenous proteins can contribute to the formation of stationary condensates in the nucleus. Furthermore, the stationary condensates would trap diffusing CRY2R in the condensate. In contrast, CRY2R condensates moved rapidly in the cytoplasm (Fig. 3). A possible explanation for this phenomenon would be the absence of scaffolding proteins such as SC35 in the cytoplasm. The moving property of CRY2R condensates in the cytoplasm may increase its usefulness as a condensation-inducing tag because it may not be necessarily extra interactions should be considered. Since the cytoplasmic localization of a neurodegenerative disease-causative aggregate-prone protein is likely to exert cytotoxicity [14,20], this moving property would be more likely to be useful for the analysis of condensation in the cytoplasm. Furthermore, some slow and stationary condensates in the cytoplasm may attach to the cellular components. This heterogeneous property of the dynamics of the condensate in the cytoplasm may be involved in the “dirty” property in the cytoplasm (e.g., heterogenic membranous organelle and cytoskeleton).

Accordingly, the subcellular compartment may importantly affect the characteristics of self-recruitment of the biomolecules in the condensates and their moving property. Interactions of condensates with specific scaffolding molecules determine a portion of the properties of the condensates (e.g., stationarity). Moreover, SICS, fluorescence intensity measurement, and single-particle tracking analysis, based on confocal fluorescence microscopy, enable clarification of the properties of the condensates in live cells.

## 5. Conclusion

We compared the assembled states and dynamics of CRY2R condensates in live cells. The CRY2R condensates showed a different property on its dynamics depending on the subcellular compartments Therefore, the environment of the subcellular com-partment may, in part, affect the characteristics of self-recruitment of bio-molecules in the condensates and their movement property in the compartment. Our findings would help to compare the subcellular compartment-dependent different characteris-tics of the condensates in the future.

## Supplementary Materials

Video S1: Movement of CRY2R condensates in the cytoplasm; Video S2: Movement of CRY2R condensates in the nucleus.

## Author Contributions

Conceptualization, A.K.; methodology, Y. H. and A.K.; validation, Y. H. and A.K.; formal analysis, Y. H. and A.K.; investigation, Y. H. and A.K.; resources, A.K.; data curation, Y. H. and A.K.; writing—original draft preparation, A.K.; writing—review and editing, Y. H. and A.K.; visualization, A.K.; supervision, A.K.; project administration, A.K.; funding acquisition, Y. H. and A.K. All authors have read and agreed to the published version of the manuscript.

## Acknowledgments

We would like to thank Ms. Asuka Murata for her technical support. We would like to thank American Journal Experts (www.aje.com) for the English language editing.

## References

1. Alberti, S.; Hyman, A.A. Biomolecular condensates at the nexus of cellular stress, protein aggregation disease and ageing. Nature reviews. Molecular cell biology 2021, 22, 196–213, doi:10.1038/s41580-020-00326-6.

2. Mittag, T.; Pappu, R.V. A conceptual framework for understanding phase separation and addressing open questions and challenges. Molecular cell 2022, 82, 2201–2214, doi:10.1016/j.molcel.2022.05.018.

3. Banani, S.F.; Lee, H.O.; Hyman, A.A.; Rosen, M.K. Biomolecular condensates: organizers of cellular biochemistry. Nature reviews. Molecular cell biology 2017, 18, 285–298, doi:10.1038/nrm.2017.7.

4. Lafontaine, D.L.J.; Riback, J.A.; Bascetin, R.; Brangwynne, C.P. The nucleolus as a multiphase liquid condensate. Nature reviews. Molecular cell biology 2021, 22, 165–182, doi:10.1038/s41580-020-0272-6.

5. Wang, B.; Zhang, L.; Dai, T.; Qin, Z.; Lu, H.; Zhang, L.; Zhou, F. Liquid-liquid phase separation in human health and diseases. Signal Transduct Target Ther 2021, 6, 290, doi:10.1038/s41392-021-00678-1.

6. Laflamme, G.; Mekhail, K. Biomolecular condensates as arbiters of biochemical reactions inside the nucleus. Commun Biol 2020, 3, 773, doi:10.1038/s42003-020-01517-9.

7. Mondal, S.; Narayan, K.; Botterbusch, S.; Powers, I.; Zheng, J.; James, H.P.; Jin, R.; Baumgart, T. Multivalent interactions between molecular components involved in fast endophilin mediated endocytosis drive protein phase separation. Nature communications 2022, 13, 5017, doi:10.1038/s41467-022-32529-0.

8. Taslimi, A.; Vrana, J.D.; Chen, D.; Borinskaya, S.; Mayer, B.J.; Kennedy, M.J.; Tucker, C.L. An optimized optogenetic clustering tool for probing protein interaction and function. Nature communications 2014, 5, 4925, doi:10.1038/ncomms5925.

9. Park, H.; Kim, N.Y.; Lee, S.; Kim, N.; Kim, J.; Heo, W.D. Optogenetic protein clustering through fluorescent protein tagging and extension of CRY2. Nature communications 2017, 8, 30, doi:10.1038/s41467-017-00060-2.

10. Mas, P.; Devlin, P.F.; Panda, S.; Kay, S.A. Functional interaction of phytochrome B and cryptochrome 2. Nature 2000, 408, 207–211, doi:10.1038/35041583.

11. Petersen, N.O.; Hoddelius, P.L.; Wiseman, P.W.; Seger, O.; Magnusson, K.E. Quantitation of membrane receptor distributions by image correlation spectroscopy: concept and application. Biophysical journal 1993, 65, 1135–1146, doi:10.1016/S0006-3495(93)81173-1.

12. Kitamura, A.; Shimizu, H.; Kinjo, M. Determination of cytoplasmic optineurin foci sizes using image correlation spectroscopy. Journal of biochemistry 2018, 164, 223–229, doi:10.1093/jb/mvy044.

13. Ershov, D.; Phan, M.S.; Pylvanainen, J.W.; Rigaud, S.U.; Le Blanc, L.; Charles-Orszag, A.; Conway, J.R.W.; Laine, R.F.; Roy, N.H.; Bonazzi, D.; et al. TrackMate 7: integrating state-of-the-art segmentation algorithms into tracking pipelines. Nature methods 2022, 19, 829–832, doi:10.1038/s41592-022-01507-1.

14. Kitamura, A.; Nakayama, Y.; Shibasaki, A.; Taki, A.; Yuno, S.; Takeda, K.; Yahara, M.; Tanabe, N.; Kinjo, M. Interaction of RNA with a C-terminal fragment of the amyotrophic lateral sclerosis-associated TDP43 reduces cytotoxicity. Scientific reports 2016, 6, 19230, doi:10.1038/srep19230.

15. Ma, L.; Guan, Z.; Wang, Q.; Yan, X.; Wang, J.; Wang, Z.; Cao, J.; Zhang, D.; Gong, X.; Yin, P. Structural insights into the photoactivation of Arabidopsis CRY2. Nat Plants 2020, 6, 1432–1438, doi:10.1038/s41477-020-00800-1.

16. Lu, M.; Williamson, N.; Mishra, A.; Michel, C.H.; Kaminski, C.F.; Tunnacliffe, A.; Kaminski Schierle, G.S. Structural progression of amyloid-beta Arctic mutant aggregation in cells revealed by multiparametric imaging. The Journal of biological chemistry 2019, 294, 1478–1487, doi:10.1074/jbc.RA118.004511.

17. Cashmore, A.R.; Jarillo, J.A.; Wu, Y.J.; Liu, D. Cryptochromes: blue light receptors for plants and animals. Science 1999, 284, 760–765, doi:10.1126/science.284.5415.760.

18. Pack, C.; Saito, K.; Tamura, M.; Kinjo, M. Microenvironment and effect of energy depletion in the nucleus analyzed by mobility of multiple oligomeric EGFPs. Biophysical journal 2006, 91, 3921–3936, doi:10.1529/biophysj.105.079467.

19. Kim, K.; Guck, J. The Relative Densities of Cytoplasm and Nuclear Compartments Are Robust against Strong Perturbation. Biophysical journal 2020, 119, 1946–1957, doi:10.1016/j.bpj.2020.08.044.

20. Asakawa, K.; Handa, H.; Kawakami, K. Optogenetic modulation of TDP-43 oligomerization accelerates ALS-related pathologies in the spinal motor neurons. Nature communications 2020, 11, 1004, doi:10.1038/s41467-020-14815-x.

